# The GLP-1 analogue, exendin-4, improves bone material properties and strength through a central relay in ovariectomized mice

**DOI:** 10.1101/2024.10.05.616809

**Authors:** Morgane Mermet, Jessica Denom, Aleksandra Mieczkowska, Emma Biggs, Fiona M. Gribble, Frank Reimann, Christophe Magnan, Celine Cruciani-Guglielmacci, Guillaume Mabilleau

## Abstract

Glucagon-like peptide-1 (GLP-1) has previously been shown to be indispensable for optimal bone strength by acting at the bone material level. However, it was not fully clear whether the effects of GLP-1 were mediated by direct or indirect actions on bone cells. In the present study, we were unable to demonstrate the expression of the GLP-1 receptor (GLP-1r) in bone tissue at the gene expression level using qPCR and in situ hybridization, or at the protein level. Furthermore, the peripheral administration of exendin-4, a specific GLP-1r agonist, in ovariectomized (OVX) BALB/c mice enhanced post-yield displacement (18%) and energy-to-fracture (24%), as well as bone volume/total volume (BV/TV) (11%), trabecular number (Tb.N) (6%), and collagen maturity (18%). These bone effects were still observed when exendin-4 was centrally administered into the lateral cerebral ventricle. On the other hand, the peripheral administration of exendin-4 coupled to bovine serum albumin, a GLP-1r agonist that cannot penetrate the brain, failed to replicate the positive effects on bone despite increased calcitonin secretion. Altogether, these data confirm that GLP-1r agonists represent an interesting approach for managing bone fragility due to ovariectomy, but also suggest that GLP-1r agonists require a central relay yet to be identified to exert positive effects on bone physiology. Further studies are needed to decipher the mechanisms of action of GLP-1 and GLP-1r agonists on bone physiology.

## 1. INTRODUCTION

Due to ageing of the population, the occurrence of bone fragility and fracture has risen significantly worldwide and will continue in the future ^(1)^. In 2019 in the European Union only, the total direct cost of fragility fractures amounted to €56.9 billion, a 64% increase compared with the figure in 2010 ^(2)^. Bone fragility is characterized by reduced bone mass, disruption of bone microarchitecture and deterioration of the quality of the bone material itself ^(3)^. Although approved anti-osteoporotic drugs increase bone mineral density, the risk of bone fracture is only reduced by 30-40% in hip and long bones in treated individuals ^(2,4-6)^. This suggests that factors beyond bone mass are important for an optimum bone strength.

Previously, we and others have highlighted the beneficial role of several gut hormones on bone physiology ^(7)^. Among gut hormones, glucagon-like peptide-1 (GLP-1) was reported to be beneficial for bone strength. Indeed, animals with a genetic impairment of the GLP-1 receptor (GLP-1r) presented with reduced bone mass, deterioration of bone microarchitecture and alterations of bone material properties, especially enzymatic collagen crosslinking, that jeopardized bone strength ^(8,9)^. In humans, GLP-1r gene polymorphisms have also been reported to impact bone mineral density and mass ^(10)^ and exogenous GLP-1 administration rapidly modulates bone remodeling ^(11-14)^. Due to the important role of GLP-1 on glucose and energy metabolism and its short half-life in the circulation, several degradation-resistant GLP-1r agonists have been developed and approved for the treatment of type 2 diabetes and more recently obesity ^(15)^. These GLP1r agonists, administered subcutaneously, have proved useful in several animal models of bone fragility to enhance bone strength ^(16-25)^. In meta-analysis of randomized clinical trials, GLP-1r agonists were reported with a neutral or slightly beneficial effects on reducing the incidence of bone fracture ^(26)^, although more recent data suggest that liraglutide and lixisenatide contribute to reduce fracture in individuals affected by type 2 diabetes mellitus ^(27)^.

However, the exact mechanism of action of GLP-1r agonist in improving bone strength remains to be elucidated. Indeed, it is not clear whether improvement of bone strength resulted from activation of skeletal GLP-1r or whether it requires extra-skeletal receptors as reported for the GIP/GIP receptor pathway ^(28)^. Furthermore, the unambiguous presence of a functional GLP-1r in bone cells remains up to now controversial due to poorly characterized cell lines, non-selective use of PCR primers or lack of sufficiently selective and commercially available antibodies ^(17,29-31)^. It has also been proposed that the bone GLP-1r would possibly be different from the conventional GLP-1r ^(32)^. Recently, despite clear effects of GLP-1 and its analogues on improving enzymatic collagen crosslinking upon peripheral administration in animals, Mieczkowska et al. were incapable of inducing a change in collagen crosslinking in vitro in osteoblast cultures ^(29)^.

The main objectives of this study were to better understand how GLP-1 controls bone strength in the ovariectomy-induced bone fragility model and to decipher whether a functional GLP-1r is expressed in bone tissue.

## 2. MATERIAL AND METHODS

### 2.1. Peptide synthesis and modification

Exendin-4 (Ex-4) and exendin-4-Cys (Ex-4-Cys), bearing a linker and an additional cysteine residue at the c-terminus, were synthesized by Genecust (Genecust, Boynes, France) to >95% purity. Peptide purity and molecular mass were determined by reverse-phase HPLC and mass spectrometry. Table 1 details the peptide sequences. To obtain exendin-4-AF647 (Ex-4-AF647), Alexa Fluor 647 maleimide (#A20347, Invitrogen, Eugene, OR) was conjugated to Ex-4-Cys according to the manufacturer’s protocol. Similarly, exendin-4-bovine serum albumin (Ex-4-BSA) was generated by conjugation of maleimide-activated BSA (#77115, ThermoScientific, Waltham, MA) with Ex-4-Cys. Exendin-4-BSA-AF647 (Ex-4-BSA-AF647) was obtained by conjugation of Ex-4-BSA with Alexa Fluor 647 NHS ester (#A20006, Invitrogen) according to the manufacturer’s protocol.

**Table 1:**
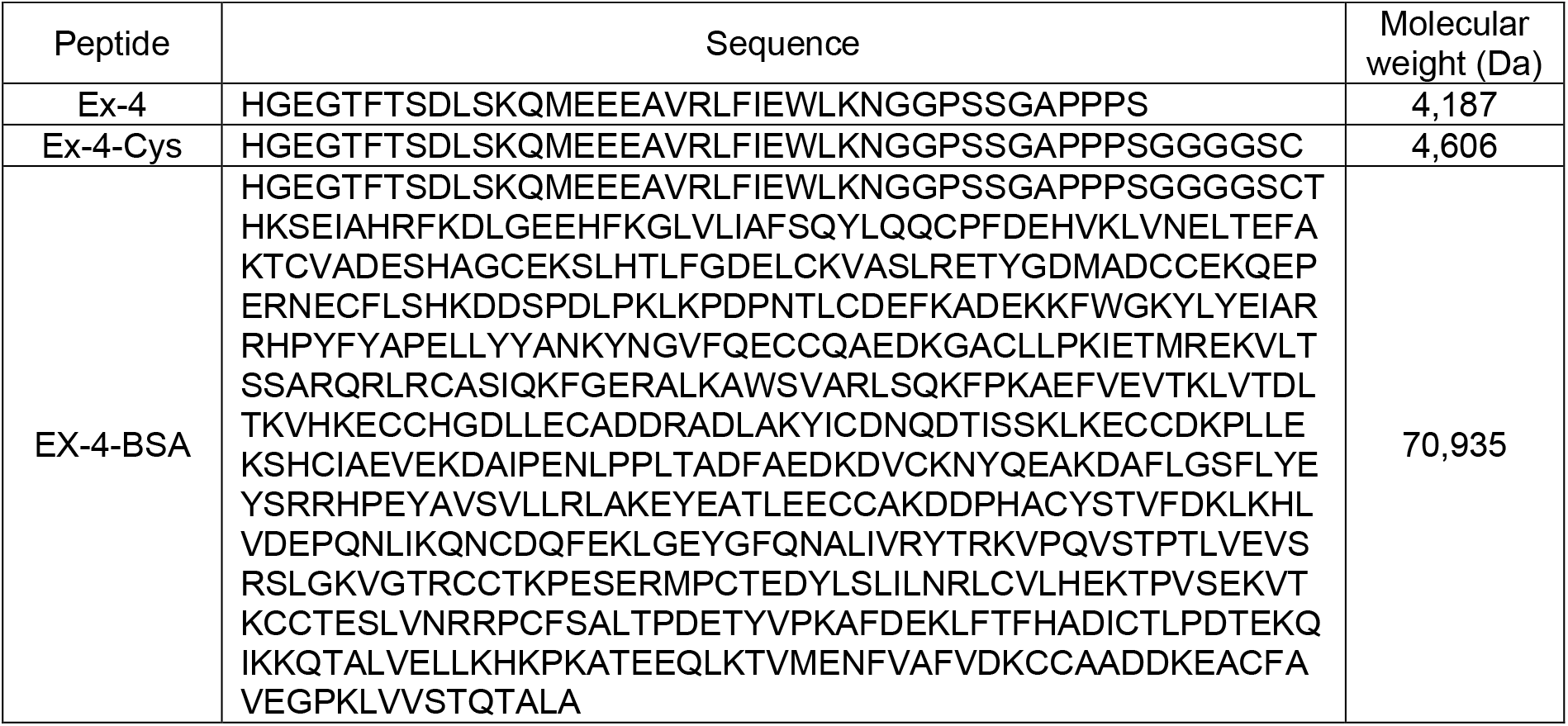
Amino acid sequences of peptides.

### 2.2. Animal models

All procedures were carried out in accordance with the European Union Directive 2010/63/EU for animal experiments and were approved by the institutional animal care and use committee of the University of Angers (CEEA-PdL N°01740.01 and 6154-201607211130415v1) and University Paris Cité (CEEA40, authorization 12952-2017022013322777v4).

#### 2.2.1. Mouse model of brain entry

Twenty 16-week-old ovariectomized female BALB/c mice (BALB/cJRj) were purchased from Janvier Laboratories (Saint-Berthevin, France). Saline (n=5), Alexa-Fluor 647 (n=5, 700 pmoles), Ex-4-AF647 (n=5, 700 pmoles) or Ex-4-BSA-AF647 (n=5, 700 pmoles) were administered subcutaneously. Thirty minutes later, the animals were sacrificed by cervical dislocation and the brains carefully dissected and frozen in liquid nitrogen. Frozen brains were powdered using a tissue biopulverizer, and proteins were extracted by incubating brain powder with RIPA buffer at 4°C for 1 hour. Samples were centrifuged at 13,000 rpm for 30 minutes at 4°C, the supernatant collected, and protein concentration determined using a BCA protein assay kit (Pierce Biotechnology, Rockford, IL). Fluorescence was measured using an excitation wavelength of 645 nm and an emission wavelength of 670 nm on a SpectraMax M2 microplate spectrofluorometer (Molecular devices, Saint-Grégoire, France).

#### 2.2.2. Mouse model of ovariectomy-induced bone fragility

Ninety-six BALB/c female mice (BALB/cJRj) were purchased from Janvier Labs (Saint-Berthevin, France). At 12 weeks of age, mice underwent sham (n=40) or bilateral (n=56) ovariectomy under general anesthesia, as previously described. ^(33)^. At 16 weeks of age, mice were given the vehicle, Ex-4 or Ex-4-BSA, for 4 weeks, either by subcutaneous (sc) or intracerebroventricular (icv) administration. For subcutaneous administration, an osmotic minipump (model 2004; Alzet, Rabalot, France) was inserted into a subcutaneous pouch on the dorsal surface of the animal under isoflurane anesthesia, and the animals received 0.05 mg/kg buprenorphine for the first 24 hours after surgery. For icv administration, animals were placed on a stereotaxic frame and a cannula was implanted in the lateral ventricle (X=-1.1mm, Y=-0.5mm and Z=-3mm positions from Bregma) under isoflurane anesthesia and received i.p. administration of xylazine 10 µg/kg. A catheter tube was connected from the cerebral perfusion cannula to an osmotic minipump (model 2004; Alzet). The minipump was inserted into a subcutaneous pocket on the animal’s dorsal surface. Correct implantation of the cerebral perfusion cannulas was verified at necropsy. Mice were chronically perfused for 28 days with vehicle, ∼700 pmol/day (sc) or 70 pmol/day (icv) Ex-4. These doses were chosen on the basis of previous publications ^(19,34,35)^. Based on the activity of Ex-4-BSA at the GLP-1r, we employed a dose of 7 nmol/day (sc).

All animals received intraperitoneal administration of calcein green (10 mg/kg) 10 days and 2 days before sacrifice. Animals were housed in social groups and maintained in a 12 h:12 h light/dark cycle. They had free access to water and food. At necropsy, blood was collected by intracardiac aspiration into EDTA-treated tubes. Blood samples were centrifuged at 13,000 g for 15 min, aliquoted and stored at −80°C until plasma levels of exendin-4 (#EK-070-94, Phoenix pharmaceuticals, Burlingame, CA), catecholamines (# BA-E-6600, Labor Diagnostika Nord GmbH, Nordhorn, Germany) and calcitonin (#NBP3-06734, Bio-Techne Ltd, Abingdon, UK) were measured according to the manufacturer’s protocol. Uteri were harvested and weighed to verify the efficacy of ovariectomy. Right femurs were wrapped in saline-soaked gauze and frozen at −20°C until use. Right tibiae were fixed in ethanol-based fixative and stored at 4°C until use.

### 2.3. High resolution X-ray microCT

MicroCT analyses were performed at the proximal metaphysis of the right tibia and at the mid-diaphysis of the right femur with a Bruker 1272 microtomograph operated at 70 kV, 140 μA, 1000 ms integration time and imaging in ethanol 70. The isotropic pixel size was fixed at 4 μm, the rotation step at 0.25° and exposure was performed with a 0.5 mm aluminum filter. Hydroxyapatite phantoms (250 mg/cm^3^ and 750 mg/cm^3^) were used for calibration. Reconstruction of 2D projections was done with the NRecon software (Version 1.6.10.2). For tibial analysis, a trabecular volume of interest was located 0.5 mm below the growth plate at the proximal end and extended 2 mm down. Trabecular bone was separated from cortical bone with an automatic contouring script in CTan software (Version 1.20.8.0). The right femur was used for cortical microarchitecture. The region analyzed (0.5 mm) was centered at the midpoint between the third trochanter and the distal growth plate. Bone was segmented from soft tissue using global thresholding set at 300 mg/cm^3^ for trabecular bone and 700 mg/cm^3^ for cortical bone. All histomorphometrical parameters were measured with the CTan software according to guidelines and nomenclature proposed by the American Society for Bone and Mineral Research ^(36)^.

### 2.4. Histomorphometrical analysis

After microCT, right tibias were embedded undecalcified in polymethylmethacrylate (pMMA) at 4°C as previously reported ^(37)^. Seven micron-thick sections were made with a sledge microtome (Polycut S, Leica, Nanterre, France) prior to histoenzymatic detection of TRAP or Goldner trichrome staining. Dynamic histomorphometry was perfomed after counterstaining the bone matrix with calcein blue. All histomorphometrical parameters were measured with an in-house written application developed in Matlab R2023b (The Mathworks, Natick, CA) according to guidelines and nomenclature proposed by the American Society for Bone and Mineral Research ^(38)^.

### 2.5. Assessment of bone strength

Whole-bone strength of right femurs was assessed by 3-point bending as described previously ^(39,40)^ and in accordance with published guidelines ^(41)^. Three-point bending strength was measured with a constant span length of 10 mm. Bones were tested in the antero-posterior axis with the posterior surface facing upward, centered on the support and the pressing force was applied vertically to the midshaft of the bone. Each bone was tested fully hydrated and at room temperature with a loading speed of 2 mm.min^−1^ until failure with a 500 N load cell on an Instron 5942 device (Instron, Elancourt, France) and the load-displacement curve was recorded at a 100 Hz rate by the Bluehill 3 software (Instron). Ultimate load, ultimate displacement, stiffness and work to fracture were calculated as indicated in ^(42)^. The yield point was determined as the point at which a regression line that represents a 10% loss in stiffness crosses the load-displacement curve. Post-yield displacement was computed as the displacement between yielding and fracture. Peak bending moment was calculated as one-half the ultimate load multiplied by one-half the span length ^(41)^. The peak bending moment is related to bone tissue material properties and bone midshaft geometry by the following equation ^(42)^:

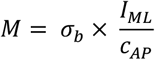

Where M is peak moment bending, σ_b_ is bone tissue material strength (TMS), I_ML_ is the moment of inertia in the medio-lateral axis, and C_AP_ is the distance from the neutral axis to bone surface in the antero-posterior direction. Differences in peak moment bending that are not explained by I_ML_/C_AP_ are caused by alterations in tissue material properties ^(43)^.

### 2.6. Bone ECM material evaluation

Distal halves of the right femur were embedded undecalcified in pMMA after dehydration and infiltration as previously reported ^(40)^. One micrometer-thick cross-section of the midshaft femur was cut with an ultramicrotome (Leica EM UC7, Leica microsystems, Nanterre, France) and deposited on BaF2 windows. Spectral analysis was performed, using a Bruker Hyperion 3000 infrared microscope coupled to a Bruker Vertex 70 spectrometer equipped with a single element mercury cadmium telluride (MCT) detector, between double calcein labels, as indicator of bone formation site. Mid-infrared spectra were recorded at a resolution of 4 cm^−1^ (Spectral range 850–2000 cm^−1^), with 32 accumulations in transmission mode. Background spectra were collected under identical conditions from the same BaF_2_ windows at the beginning and end of each experiment to ensure instrument stability. Post-processing was performed using Matlab R2023b (The Mathworks, Natick, CA) and included Mie scattering correction, pMMA subtraction, normalization and denoising (Savitzky-Golay algorithm, degree 2, span 9) prior to second derivative spectroscopy and curve fitting routines as previously reported in detail ^(44)^. A signal-to-noise ratio was computed in the region 1850-2000 cm^−1^ to ensure proper denoising and quality of FTIR spectra for subsequent second derivative and curve fitting. Bone ECM material parameters were: Phosphate/Amide ratio (area ratio of v1,v3 phosphate and amide I); mineral crystallinity/maturity (XST, area ratio of subbands located at subbands ∼1030 cm^−1^ and ∼1020 cm^−1^) ^(45)^, carbonate/phosphate ratio (area ratio of v2 carbonate and v1,v3 phosphate) ^(45)^, and collagen maturity (intensity ratio of subbands located at 1660 cm^−1^/1690 cm^−1^) ^(44)^.

### 2.7. Expression of Glp1r

Femurs from 12-week-old female BALB/c (BALB/cJRj) mice were rapidly dissected, cleared of soft tissue before the distal ends were cut. Bones were centrifuged for 20 seconds at 16,000 g at 4°C, as previously described in detail ^(46)^, to remove bone marrow. Bone tissue was frozen in RNA later in liquid nitrogen and stored at −80°C until use. Hypothalamus, pancreas and sinus node region were used as positive controls and frozen in RNA later. Liver and skeletal muscle were used as negative controls and processed as described above. After thawing, RNA was extracted by grinding tissue in Nucleozol (Macherey-Nagel, Hoerdt, France) and purifying total RNA with Nucleospin RNA columns (Macherey-Nagel) according to the manufacturer’s recommendations. Total RNA was reverse transcribed using the maxima first strand cDNA synthesis kit (Thermofisher scientific, Carlsbad, USA). Real-time quantitative polymerase chain reaction (qPCR) was performed using a Bio-Rad CFX 96 system and TaqMan gene expression assays (Thermofisher Scientific). Taqman assay ID # Mm00445292_m1 was used to assess the relative expression of mouse Glp1r versus mouse B2m (Taqman assay ID # Mm00437762_m1).

In addition, femurs from 5-week-old female BALB/c mice (BALB/cJRj) were rapidly dissected and fixed in formalin for 16 h before being decalcified in autoclaved 10% EDTA and embedded in paraffin following standard protocols. Tissues were cut into 5 µm sections and mounted on Superfrost plus gold glass slides (Thermofisher Scientific). In situ hybridization (ISH) was performed using the RNAscope v2 multiplex fluorescent reagent kit (Advanced Cell Diagnostics Inc., Newark, CA) and specific murine Glp1r (Assay No.: Mm-Glp1r #418851), murine Col1a1 (Assay No.: Mm-Col1a1 #319371) hybridization probes according to the manufacturer’s instructions and counterstained with DAPI. Slides were imaged with a Leica SP8 confocal microscope (Leica microsystems). Murine pancreases served as positive controls and were simultaneously processed as described above for bone samples and analyzed by RNAscope. In addition, femurs and pancreases were isolated from 13-week-old GLP-1R-Cre-eYFP mice, expressing cytosolic fluorescent eYFP only in cells expressing GLP-1R ^(47)^. Briefly, femurs and pancreases were fixed in 2.5% formaldehyde in 1X PBS, decalcified in 10% EDTA and embedded in paraffin. Five µm-thick sections were cut and mounted on Superfrost plus Gold glass slides (Thermofisher scientific). eYFP expression was assessed by immunohistochemistry (IHC) using anti-YFP (1/200 dilution, Abcam # Ab5450) incubated overnight at 4°C followed by an AlexaFluor 647-coupled secondary antibody (Thermofisher Scientific). Slides were counterstained with DAPI and imaged with a Leica SP8 confocal microscope (Leica microsystems).

### 2.8. Receptor binding and cAMP assays

The sequence encoding murine Glp1r was cloned into the pcDNA3.1(+) vector between the NheI and EcoRI sites of multiple cloning sites. CHO-K1 cells were grown and expanded at a 1:5 ratio in propagation medium containing F-12 supplemented with 10% FBS, 100 U/ml penicillin and 100 mg/ml streptomycin in a humidified atmosphere enriched with 5% CO2 at 37°C. CHO-K1 cells were transfected with the plasmid encoding the GLP-1 receptor using Invitrogen™ Lipofectamine™ 3000 as recommended by the manufacturer. For binding assay, 24h post-transfection, cells were detached and plated at a density of 6×104 cells/cm2 in black 96 well plates with clear bottom (Ibidi GmbH, Martinsried, Germany). After 24h, cells were exposed to various concentrations of Ex-4 or Ex-4-BSA in the presence of 10-6M Fam-[D-Ala2]-GLP-1 in α-MEM supplemented with 0.1% bovine serum albumin. Equilibrium binding was achieved overnight at 37°C. Cells were then washed twice with PBS and solubilized in 0.1M NaOH. Fluorescence was read with on a SpectraMax M2 microplate spectrofluorometer with excitation and emission wavelength set up at 490 nm and 525 nm respectively.

For activation assay, plasmid encoding the mGLP-1r and the Epac-S-H74 probe, a validated cAMP FRET biosensor ^(48)^, were transfected with Invitrogen™ lipofectamine™ 3000 as described above. Forty-eight hours later, transfected cells were incubated in HEPES buffered saline in the presence of Ex-4-based peptides for 30 minutes. Donor excitation was made at 460 nm, donor emission was collected at 480 nm and acceptor emission at 560 nm with a SpectraMax M2 microplate spectrofluorometer. FRET was expressed as the ratio between donor and acceptor signals. The FRET ratio was standardized at 1 with vehicle. An increase in FRET ratio suggests an augmentation in intracellular cAMP levels and hence receptor activation.

### 2.9. Statistical analysis

Statistical analyses were performed with GraphPad Prism 8.0 (GraphPad Software, La Jolla, CA, USA). The Brown-Forsythe test for equality of variance was applied to verify that normality was respected. An ordinary one-way ANOVA with multiple Tukey post hoc comparisons was performed. Outliers were identified using the ROUT method with a threshold of 1%. A linear regression between collagen maturity and tissue material strength was performed and Pearson’s correlation coefficient (r) calculated. Binding at the mGLP-1r was achieved by non-linear regression using the one-site Fit log IC50 model. Unless otherwise indicated, data are presented as mean ± SD. Differences at p<0.05 were considered significant.

## 3. RESULTS

### 3.1. Subcutaneous administration of exendin-4 improves bone strength and bone material properties in OVX animal

Ex-4 was administered peripherally via subcutaneous implantation of osmotic minipumps. As expected, ovariectomy resulted in a significant deterioration in the mechanical response of long bones, represented by decreases in ultimate load (−15%, p<0.001), stiffness (−18%, p=0.004), post-yield displacement (−18%, p=0.005) and energy-to-fracture (−24%, p=0.003) (Figures 1A-D). Subcutaneous administration of Ex-4 improved plastic deformation of the femurs, as shown by significant increases in post-yield displacement (18%, p=0.025) and energy-to-fracture (24%, p=0.020). On the other hand, subcutaneous administration of Ex-4 did not prevent the impact of ovariectomy on ultimate load (p=0.817) and stiffness (p=0.984).

**Figure 1:**
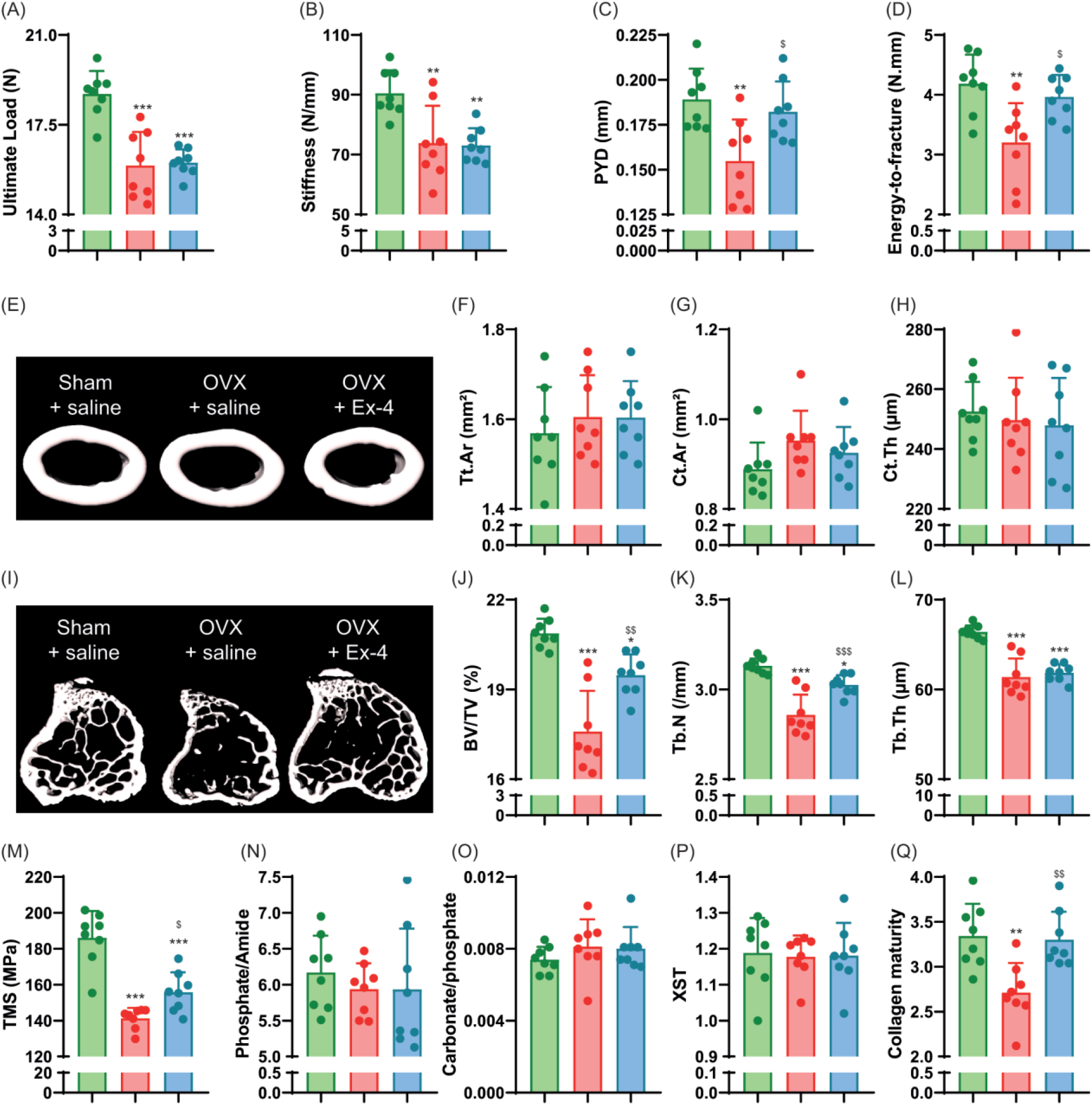
Investigation of bone phenotype following subcutaneous administration of exendin-4. (A-D) Mechanical properties of the femur and (E-H) cortical bone microstructure were investigated at the midshaft femur by 3-point bending and microCT. (I-L) Trabecular microstructure was investigated at the proximal tibia metaphysis. (M-Q) Bone material strength and properties were investigated at the midshaft femur. In panels A-Q, Green bars = Sham + vehicle, red bars = OVX + vehicle and blue bars = OVX + Ex-4. Statistical analyses were performed with one way ANOVA with Tukey’s multiple comparisons test. *: p<0.05; **: p<0.01; ***: p<0.001 vs. Sham+Vehicle. ^$^: p<0.05; ^$$^: p<0.01; ^$$$^: p<0.001 vs. OVX+Vehicle

To better understand whether the positive effects of Ex-4 were related to changes in femur microstructure, we assessed bone microarchitecture at the mid-diaphysis level (Figures 1E-H). Interestingly, OVX animals showed no significant changes in cortical bone microstructure compared with sham animals. Similarly, subcutaneous administration of Ex-4 did not alter cortical bone microstructure in OVX animals despite positive effects on biomechanics. Trabecular microarchitecture was assessed at the proximal metaphysis of the tibia (Figure 1I-1L). As expected, OVX led to decreases in bone mass represented by lower BV/TV (−16%, p<0.001), Tb.N (−9%, p<0.001) and Tb.Th (−8%, p<0.001). Interestingly, subcutaneous administration of Ex-4 resulted in higher BV/TV (11%, p=0.002) and Tb.N (6%, p<0.001). Histological evaluation revealed no effects of ovariectomy on dynamic morphometry or osteoid parameters (Table 2).

**Table 2:**
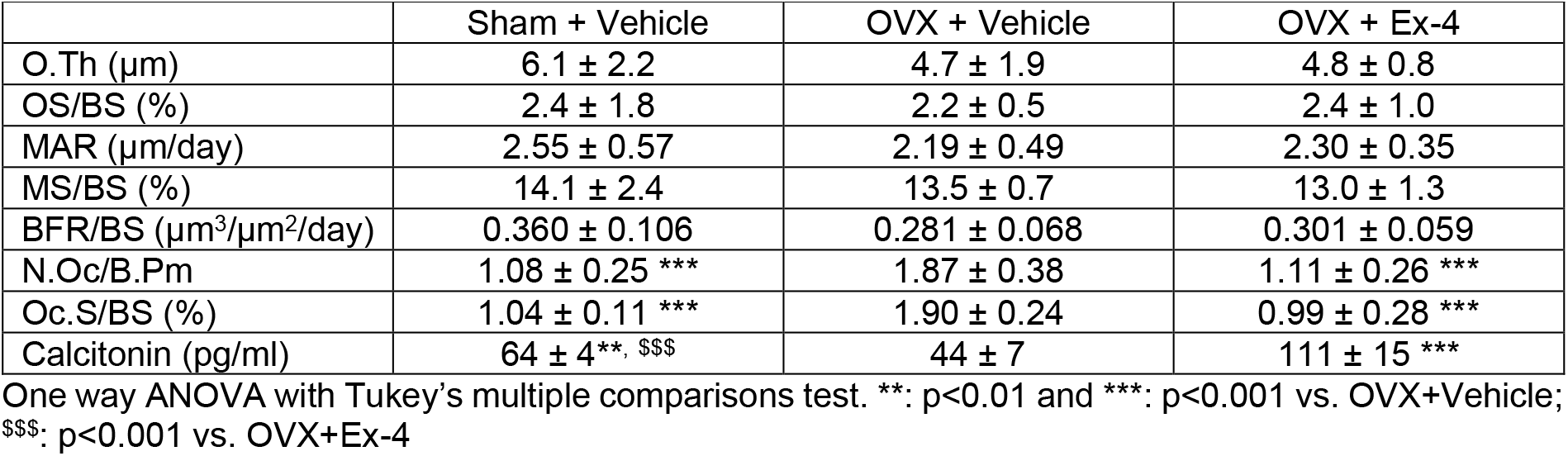
Bone histomorphometry at the proximal tibia metaphysis and plasma levels of calcitonin after subcutaneous administration of Ex-4.

On the other hand, the number of osteoclasts and osteoclast surfaces were significantly increased in OVX+vehicle animals (76%, p<0.001) and restored after subcutaneous administration of Ex-4 (−41%, p<0.001). Plasma circulating levels of calcitonin were also significantly reduced in vehicle-treated ovariectomized animals (−31%, p=0.001) and significantly increased in OVX animals treated with subcutaneous Ex-4 (152%, p<0.001) (Table 2)

We then calculated tissue material strength to determine whether the mechanical response was related to changes in bone material. Interestingly, OVX animals showed a significant reduction in tissue material strength compared to sham animals (−24%, p<0.001) which was partially reversed by Ex-4 administration (11%, p=0.043), suggesting that the main effect of Ex-4 on cortical bone strength was related to changes in bone material (Figure 1M). At the material level, the bone mineral was not affected by OVX or subcutaneous administration of Ex-4 (Figures 1N-P). In contrast, collagen maturity, representing the ratio of pyridinoline to dihydroxylysinonorleucine, was significantly lower in vehicle-treated ovariectomized animals (−19%, p=0.003) and significantly improved by subcutaneous administration of Ex-4 (18%, p=0.005) (Figure 1Q). Interestingly, collagen maturity correlated linearly with tissue material strength (r = 0.50, p=0.012) and energy-to-fracture (r =0.63, p =0.001).

### 3.2. GLP-1r receptor is not expressed in bone

As exendin-4 exerts positive effects on the properties of bone material and in particular on collagen maturity, we wanted to determine whether Ex-4 acted directly on osteoblasts. As expected, Glp1r expression was found in pancreas, heart, lung and hypothalamus (positive controls) and absent in liver and skeletal muscle (negative controls). Interestingly, we were unable to detect Glp1r expression in bone by qPCR (Figure 2A). We also explored Glp1r expression by in situ hybridization with the RNAscope assay. Although Glp1r expression was found in pancreatic islets, we failed to demonstrate the presence of Glp1r expression in bone cells or bone marrow (Figure 2B). We also investigated the presence of YFP-positive cells in the pancreas and long bones of GLP-1R-CRE-eYFP mice (Figure 2C). Again, we demonstrated the presence of YFP-positive cells in pancreatic islets but failed to demonstrate the presence of YFP-positive cells in bone marrow or bone tissue. However, several articular chondrocytes were YFP-positive, suggesting that during articular chondrocyte differentiation, the Glp1r promoter was activated and led to transcriptional and translational expression of YFP. Overall, these data suggest that the positive effects of subcutaneous Ex-4 administration on bone mechanics and fracture resistance are indirect.

**Figure 2:**
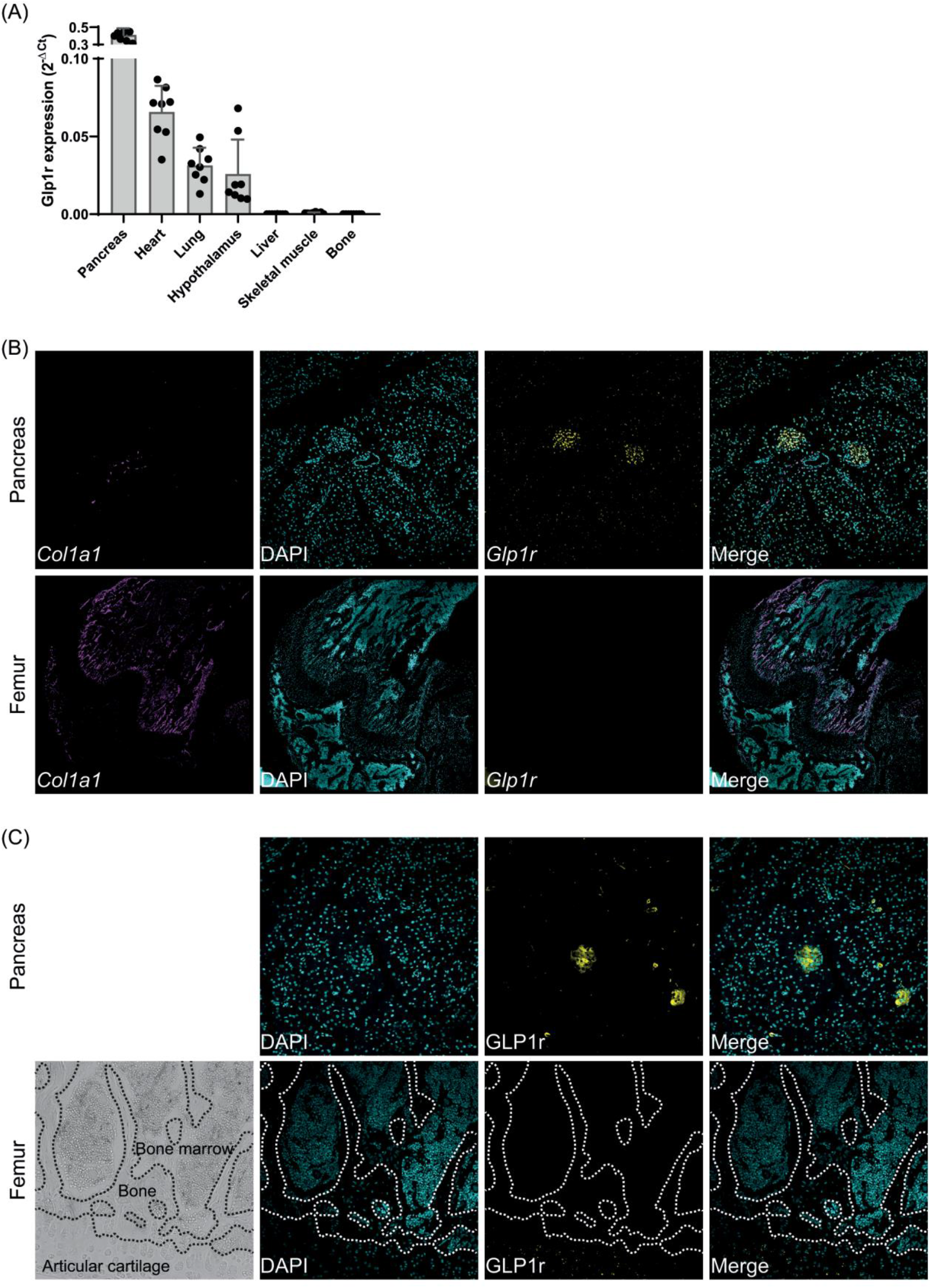
Expression of Glp1r in bone. Glp1r expression was investigated by (A) qPCR or (B) in situ hybridization using the RNAscope technology. Pancreases, hearts, lung and hypothalamus were used as positive controls. Liver and skeletal muscle were used as negative controls. (C) Expression of GLP-1r was also investigated at the protein level in pancreas and bone extracted from GLP-1R-CRE-eYFP mice. Dashed lines represent the contour of bone trabecula.

### 3.3. Intracerebroventricular administration of exendin-4 improves bone strength and bone material properties in OVX animal

As liraglutide, another GLP-1r analogue was able to enhance bone resistance to fracture in a model of streptozotocin-induced type 1 diabetes mellitus^(20)^ characterized by chemical destruction of pancreatic islets, we hypothesized that Ex-4 mediates its effects through activation of receptors in other tissues, including the CNS. We first examined whether Ex-4 could enter the brain by using Ex-4-AF647. Within 30 minutes of subcutaneous administration, Ex-4-AF647 was detected in brain extracts, suggesting that not only Ex-4-AF647, but also possibly Ex-4, were able to access the CNS at least in circumventricular organs and adjacent brain nuclei (Figure 3A). We then administered saline or Ex-4 intracerebroventricularly (icv) via a cannula inserted into the lateral ventricle and connected it to a peripheral osmotic minipump in ovariectomized animals. Ovariectomized mice treated with icv saline exhibited a significant reduction in ultimate load (−19%, p<0.001), stiffness (−16%, p<0.001), post-yield displacement (−16%, p<0.001) and energy-to-fracture (−15%, p<0.001) compared to sham control animals (Figures 3B-E). Interestingly, icv administration of Ex-4 resulted in a significant improvement in ultimate load (7%, p=0.010) and plastic deformation, represented by higher post-yield displacement (8%, p<0.001) and energy-to-fracture (8%, p<0.001). We then tested whether the positive effects on biomechanics following icv administration of Ex-4 were due to changes in cortical bone microstructure. However, here again and similarly to what was found for subcutaneous administration of Ex-4, we were unable to demonstrate significant changes in cortical bone microstructure (Figures 3F-I). The microstructure of the trabecular bone was studied in the proximal metaphysis of the tibia (Figs. 3J-M). Interestingly, vehicle-treated OVX animals showed a decrease in BV/TV (−10%, p<0.001), Tb.N (−5%, p<0.001) and Tb.Th (−5%, p<0.001), confirming the results observed in the above-mentioned subcutaneous study. Intracerebroventricular administration of Ex-4 reversed alterations in BV/TV (+6%, p=0.005) and Tb.N (4%, p<0.001).

**Figure 3:**
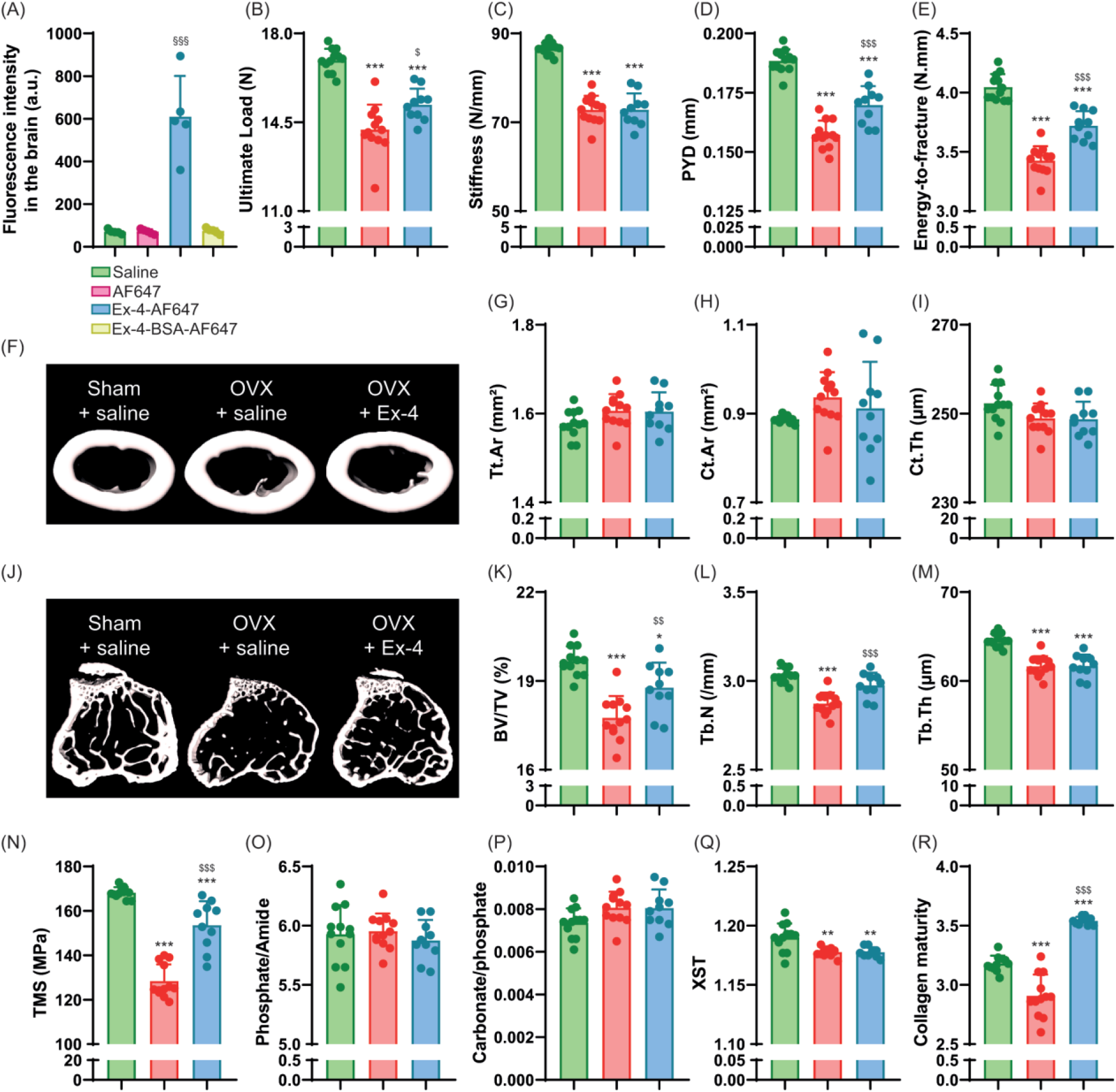
Investigation of bone phenotype following intracerebroventricular administration of exendin-4. **(A)** Crossing of the blood-brain barrier was investigated by using fluorescently labelled Ex-4. **(B-E)** Mechanical properties and (F-I) cortical bone microstructure were investigated at the midshaft femur by 3-point bending and microCT. (J-L) Trabecular microstructure was investigated at the proximal tibia metaphysis by microCT. (M-Q) Bone material strength and properties were investigated at the midshaft femur. In panels B-R, Green bars = Sham + vehicle, red bars = OVX + vehicle and blue bars = OVX + Ex-4. Statistical analyses were performed with one way ANOVA with Tukey’s multiple comparisons test. ^§§§^: p<0.001 vs. saline; *: p<0.05; **: p<0.01; ***: p<0.001 vs. Sham+Vehicle. ^$^: p<0.05; ^$$^: p<0.01; ^$$$^: p<0.001 vs. OVX+Vehicle

Histological analyses again revealed that the main effects of OVX concerned osteoclast numbers (73%, p=0.003) and osteoclast surfaces (107%, p<0.001) (Table 3). Intracerebroventricular administration of Ex-4 reduced the number of osteoclasts (−51%, p<0.001) and osteoclast surfaces (−45%, p<0.001), but had no effects on circulating calcitonin levels.

**Table 3:**
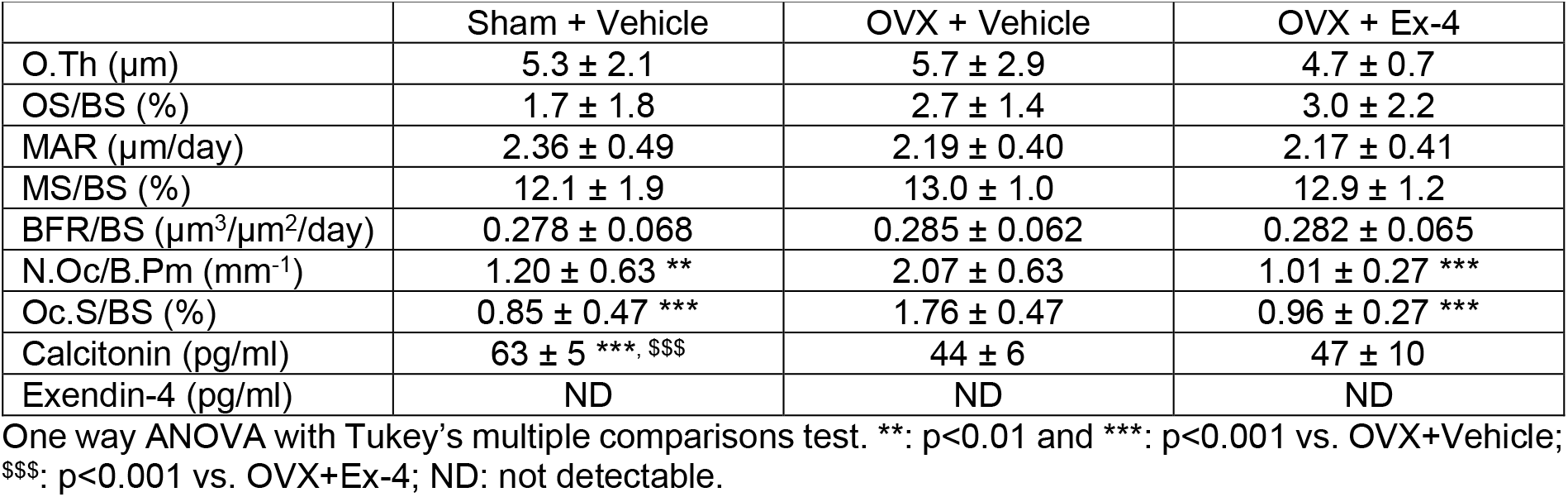
Bone histomorphometry at the proximal tibia metaphysis and plasma levels of calcitonin and exendin-4 after intracerebroventricular administration of Ex-4.

We then calculated tissue material strength and found that ovariectomized animals treated with icv saline had significantly lower tissue material strength than sham animals (−24%, p<0.001) (Figure 3N). Interestingly, ovariectomized animals treated with icv Ex-4 showed a significant improvement in this parameter (9%, p=0.005), suggesting an effect of Ex-4 on bone material properties via a central relay. At material level, vehicle-treated OVX animals showed normal mineral properties (Figures 3O-R). In contrast, collagen maturity was significantly reduced in OVX animals (−8%, p<0.001). Intracerebroventricular administration of Ex-4 in OVX animals improved collagen maturity (22%, p<0.001). Interestingly, collagen maturity correlated linearly with tissue strength (r=0.70, p<0.0001) and energy-to-fracture (r=0.39, p=0.021).

### 3.4. Subcutaneous administration of exendin-4-BSA failed to improve bone strength and bone material properties in OVX animal

To verify that Ex-4 mode of action on bone biomechanics and material properties involves activation of a central relay, we designed the Ex-4-BSA agonist, corresponding to Ex-4 coupled to bovine serum albumin. We first verified that Ex-4-BSA was unable to enter the brain and as expected, Ex-4-BSA-AF647 was not found in brain extracts (Figure 3A). Because of potential steric interference of the BSA moiety with GLP-1r interaction we then assessed whether Ex-4-BSA could still bind to murine GLP-1r (Figure 4A).

**Figure 4:**
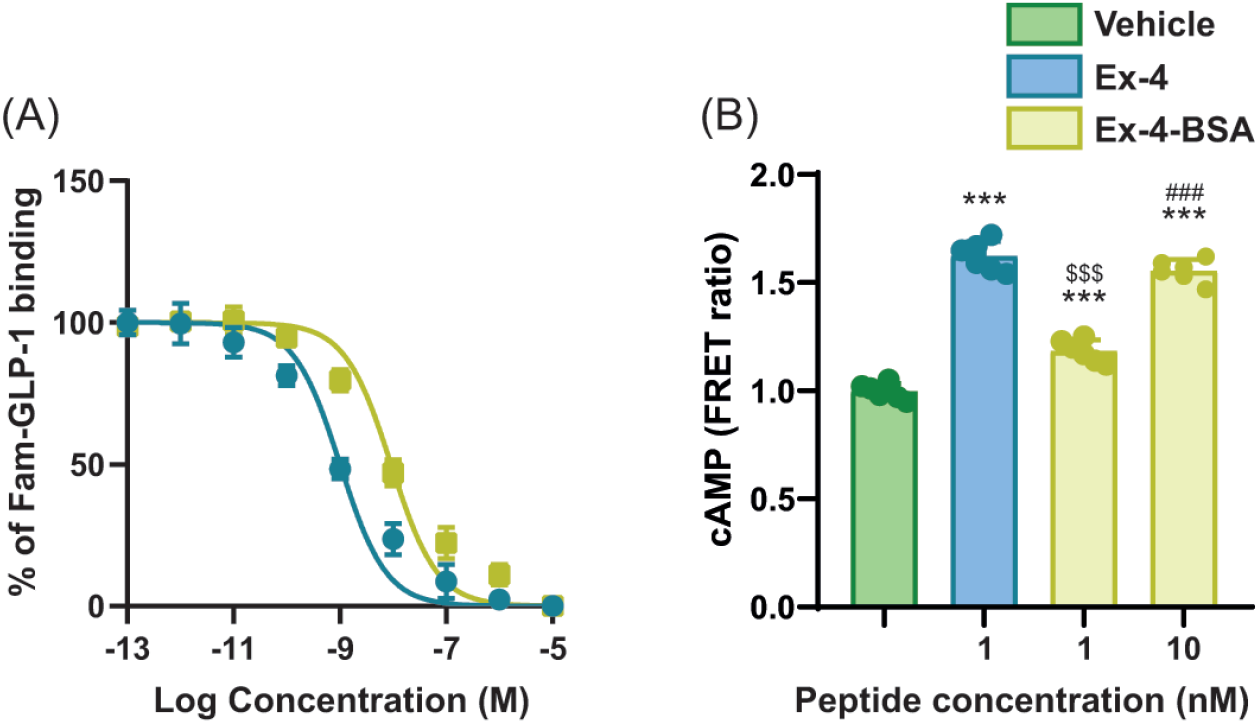
Binding to and activation of the mGLP-1r by Ex-4-BSA. (A) Competitive binding assay of Ex-4-BSA in mGLP-1r-transfected CHO-K1 cells. (B) Cyclic AMP production was detected by FRET using the H74 cAMP biosensor probe in mGLP-1r-transfected CHO-K1 cells. Production of cAMP is seen by increasing FRET ratio. Statistical analyses were performed with one way ANOVA with Tukey’s multiple comparisons test. ***: p<0.001 vs. Vehicle; ^$$$^: p<0.001 vs. Ex-4 and ^###^: p<0.001 vs. 1 nM Ex-4-BSA.

Compared with Ex-4, Ex-4-BSA showed a ∼1 log difference in binding activity (EC50 = 9.8 × 10^−9^ M ± 3.5 × 10^−9^ M vs. 1.1 × 10^−9^ M ± 0.4 × 10^−9^ M, p<0.001) (Figure 4A). We next ascertained whether Ex-4-BSA was also capable of activating the mGLP-1r (Figure 4B). Ex-4-BSA was administered in cultures of CHO-K1 cells transfected with the mGLP-1r and the H74 cAMP biosensor. As compared with Ex-4, Ex-4-BSA was capable of inducing the intracellular production of cAMP but with a concentration ∼10 times higher, suggesting that it could still activates the mGLP-1r (Figure 4B). Ex-4-BSA was then subcutaneously administered with an osmotic minipump into ovariectomized mice at a concentration of 7 nmol/day. Interestingly, Ex-4-BSA did not improve any of the biomechanical parameters, that were all affected by ovariectomy (Figures 5A-D). Similarly, Ex-4-BSA had no effect on cortical and trabecular bone microarchitectures (Figures 5E-L).

**Figure 5:**
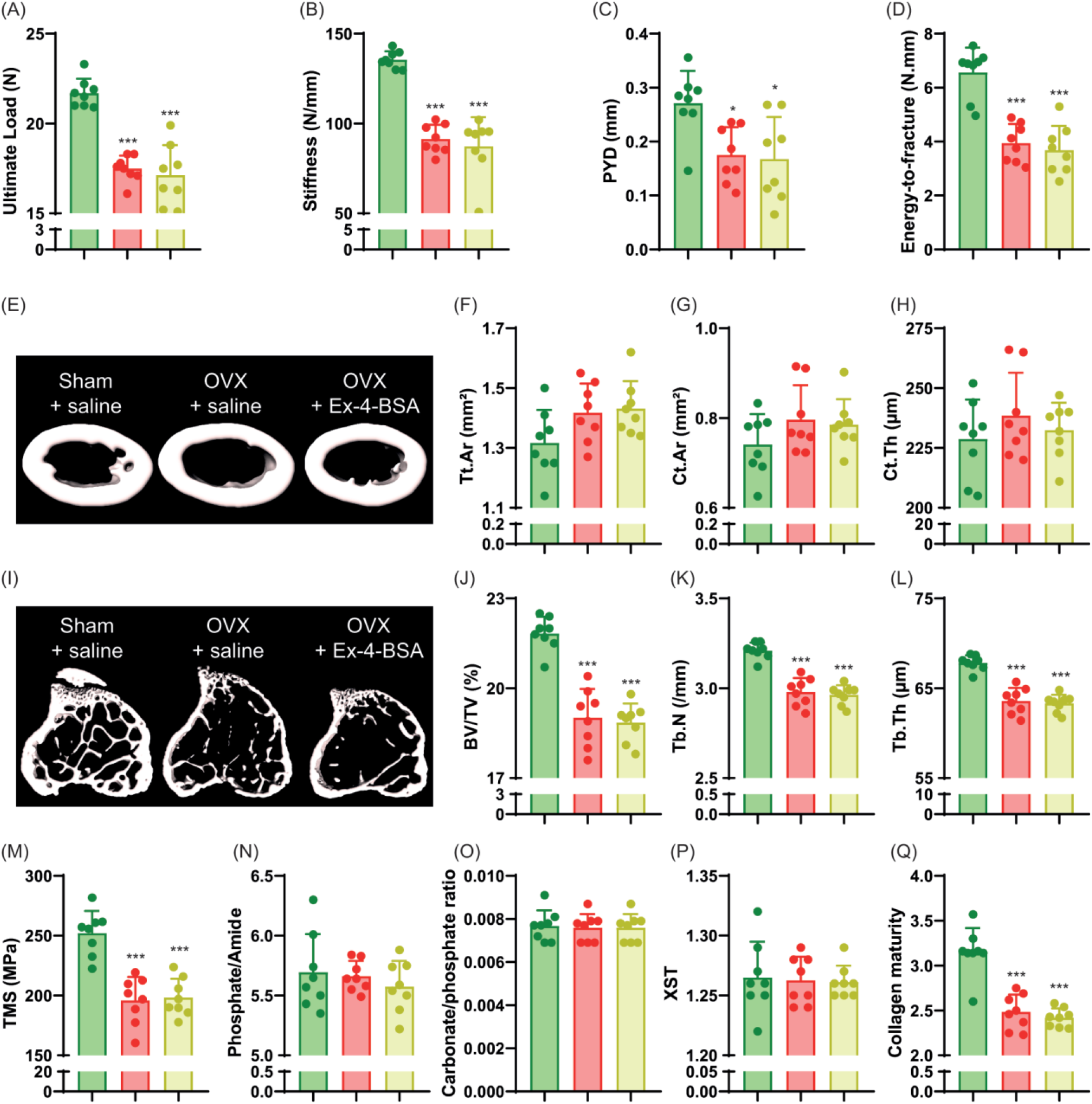
Investigation of bone phenotype following subcutaneous administration of exendin-4-BSA. Mechanical properties of the femur (A-D) and cortical microstructure (E-H) were investigated at the midshaft femur. (I-L) Trabecular microstructure was investigated at the proximal tibia metaphysis. (M-Q) Bone material strength and properties were investigated at the midshaft femur. In panel A-Q, Green bars = Sham + vehicle, red bars = OVX + vehicle and yellow bars = OVX + Ex-4-BSA. Statistical analyses were performed with one way ANOVA with Tukey’s multiple comparisons test. *: p<0.05; **: p<0.01; ***: p<0.001 vs. Sham+Vehicle. ^$^: p<0.05; ^$$^: p<0.01; ^$$$^: p<0.001 vs. OVX+Vehicle

Histological analyses failed to evidence any effects of Ex-4-BSA on the number of osteoclasts or osteoclast surfaces. However, Ex-4-BSA induced an increase in circulating calcitonin levels (152%, p<0.001) suggesting that Ex-4-BSA was active at peripheral sites (Table 4). Subcutaneous administration of Ex-4-BSA did not improve tissue material strength (p=0.962) (Figure 5M). Furthermore, at the material level, Ex-4-BSA did not modify collagen maturity in OVX animals (p=0.802) (Figure 5Q).

**Table 4:**
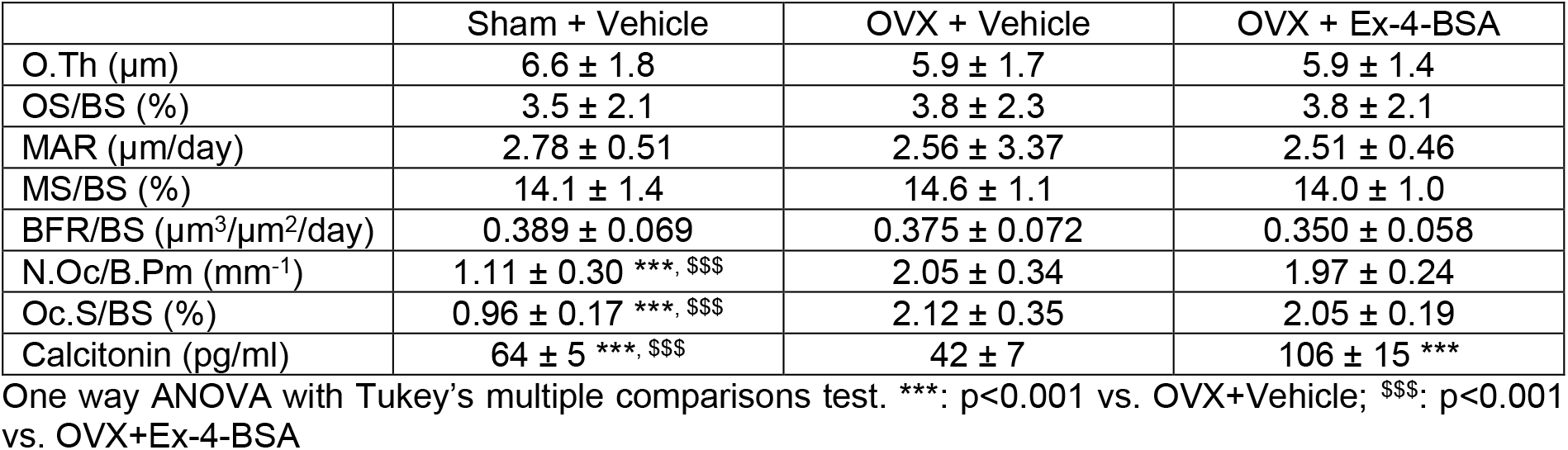
Bone histomorphometry at the proximal tibia metaphysis and plasma levels of calcitonin after subcutaneous administration of Ex-4-BSA.

## 4. DISCUSSION

Bone fragility is on the rise worldwide, and new strategies are needed to better manage patients at risk of bone fracture. Among these, the repurposing of GLP-1r agonists, approved for the treatment of type 2 diabetes and obesity, could represent a valuable solution.

In the present study, we unambiguously demonstrated that subcutaneous or intracerebroventricular administration of the GLP-1r agonist, exendin-4, was capable of improving not only the microstructure of trabecular bone, but also the strength of cortical bone, through increased collagen maturity. We have also shown that Ex-4 enters the brain. These results were not observed in animals treated with Ex-4-BSA, suggesting that the action of Ex-4 is central. Consistent with a central relay, we failed to demonstrate GLP-1r expression in bone tissue. GLP-1r expression has previously been reported in the arterial walls of the kidneys and lungs, cardiac myocytes of the sinus node, gastric antrum and pylorus, enteric neurons, vagal and dorsal root ganglia, β-cells of the pancreas and c-cells of the thyroid gland ^(47,49)^. Several central regions, including the olfactory bulb, amygdala, preoptic gland and brain, amygdala, preoptic area, nucleus accumbens, hypothalamic arcuate nucleus, paraventricular nucleus, dorsomedial nucleus, lateral hypothalamic nucleus, supraoptic nucleus, postrema area, nucleus tractus solitarius and lateral reticular nucleus express the GLP-1r ^(47,49-52)^. It should be noted that we observed a slight but significant expression of YFP protein in articular chondrocytes from GLP-1r-Cre-eYFP mice, suggesting that the Glp1r promoter was activated during articular chondrocyte differentiation. However, why *Glp1r* transcripts were not revealed by ISH in articular chondrocytes is currently unclear. Nevertheless, GLP-1r expression by articular cartilage has also been reported previously by Meurot et al.^(53)^. Pancreatic GLP-1r activation is unlikely to be responsible for the observed increase in collagen maturity and bone strength upon Ex-4 administration, as animals with significant pancreatic islet destruction by streptozotocin still showed the same type of response. ^(20)^.

Global Glp1r KO animals show alterations in bone strength, reduced trabecular bone mass and material properties, suggesting GLP-1 control of these bone functions ^(8,9)^. These observations formed the basis for investigating the potential utility of a GLP-1r agonist for the management of bone fragility. Ex-4 has already been administered subcutaneously in preclinical models of bone fragility, including the OVX model used in the present study. Previous studies have reported positive effects of Ex-4 that increased bone mineral density, trabecular microstructure and circulating calcitonin levels and reduced osteoclast parameters, resulting in greater bone strength ^(16,19,24)^. Our study recapitulates these findings, but more importantly highlights that although Ex-4 can stimulate calcitonin secretion by the thyroid gland in rodents ^(54)^, the effects on trabecular and cortical bone microstructure, material properties and strength are due to activation of a central relay. Furthermore, despite a significant rise in circulating calcitonin levels in response to Ex-4-BSA, this GLP-1r agonist is unable to affect bone physiology, suggesting that the positive effects of Ex-4 on bone may be independent of calcitonin secretion in rodents.

Interestingly, intracerebroventricular administration of Ex-4 recapitulated the greater collagen maturity observed after subcutaneous administration, suggesting that central GLP-1r is required to modulate the quality of the bone material component. This hypothesis is reinforced by the lack of effect of Ex-4-BSA, which does not enter the brain, on bone material properties. This result is in line with a previous in vitro study which showed no direct positive effects of GLP-1 on collagen maturity ^(29)^. Bone fragility is governed by the triad of bone mass - bone microstructure - bone material properties. The idea that bone mass can be controlled by central relays is not new, and dates back to the early 2000s with the pioneering work of the Karsenty lab ^(55)^ which has now been replicated and extended by several investigators worldwide and has been the subject of several reviews over the last decade ^(56-60)^. However, to our knowledge, the present study is the first report of bone quality control by a central relay. Of course, further work is needed to decipher which central region is involved in this process and how centrally generated signals can be projected to the skeleton.

Furthermore, although the GLP-1 sequence is conserved between rodents and humans, the tissue distribution of GLP-1r differs between the two species, with the example of Glp1r expression in thyroid C cells in rodents but not in non-human primates or humans. ^(54)^. In the present study, Ex-4-BSA failed to exhibit significant improvement in collagen maturity and hence bone strength, despite increased secretion of calcitonin in animals and activity at the murine GLP-1r in transfected CHO cells. This suggests that the beneficial effects of Ex-4, and possibly GLP-1, on collagen maturity and bone strength is not mediated by either calcitonin secretion or peripheral action, but rather by activation of a central relay. With respect to the lack of expression of *Glp1r* in human thyroid C-cells, it would be interesting to investigate whether the central action of Ex-4 is preserved in humans and could represent a viable option to treat bone fragility due to poor bone material quality.

A limited number of studies have investigated whether endogenous GLP-1 or GLP-1r agonists were capable of entering the brain. ^125^I-labeled Ex-4 has been shown to rapidly enter the brain in a non-saturable manner at doses below 30 nmol/kg ^(61,62)^. Furthermore, CNS access of intravenously infused radiolabeled [Ser8]GLP-1, a stable analogue of GLP-1, is not inhibited by excess doses of unlabeled [Ser8]GLP-1 or by the GLP-1r antagonist exendin (9-39) in mice ^(63)^. Taken altogether, these studies might support a brain entry through adsorptive transcytosis across brain endothelial cells. The lack of active receptor-mediated transport of GLP-1 is further supported by recent transcriptomic data failing to undoubtedly show the expression of Glp1r in hypothalamic endothelial cells or cells of the neurovascular unit ^(64)^. Brain entry is not limited to Ex-4 and lipidation of GLP-1r agonists does not seem to be a problem for brain entry as lipidated Ex-4, liraglutide and semaglutide have also been reported to enter the brain parenchyma after peripheral administration ^(65-67)^.

In conclusion, the present study identified a new mode of action of exendin-4 through a central relay in order to induce significant improvement in bone strength through action at the bone material level and collagen maturity. Whether such mechanism of action is preserved in humans remains to be elucidated.

## 5. AUTHOR CONTRIBUTION

**Morgane Mermet:** Investigation, Formal analysis, Writing – Review & Editing; **Jessica Denom:** Investigation, Formal analysis, Writing – Review & Editing; **Aleksandra Mieczkowska:** Investigation, Formal analysis, Writing - Review & Editing; **Emma Biggs:** Resources, Writing - Review & Editing, **Fiona Gribble:** Resources, Writing - Review & Editing; **Frank Reimann:** Resources, Writing - Review & Editing; **Christophe Magnan:** Conceptualization, Resources, Writing - Review & Editing; **Céline Cruciani-Guglielmacci:** Conceptualization, Resources, Writing - Review & Editing, Supervision; **Guillaume Mabilleau:** Conceptualization, Investigation, Formal analysis, Writing – original draft, Supervision, Funding acquisition, Data curation

## 6. ACKNOWLEDGEMENTS

The authors thank the SCAHU and SCIAM platforms – SFR ICAT 4208 – University of Angers and the Buffon Animal Facility– Université Paris Cité for support with animal housing and care, and confocal microscopy. This work was supported by a grant from Société Française de Rhumatologie (#2020-2650). Work in the Reimann/Gribble laboratories is supported by Wellcome (220271/Z/20/Z) and MRC-UK (MRC_MC_UU_12012/3).

## 8. DATA AVAILABILITY STATEMENT

The data that support the findings of this study are available from the corresponding author upon reasonable request.

